# Connected Chromatin Amplifies Acetylation-modulated Nucleosome Interactions

**DOI:** 10.1101/2024.10.11.617935

**Authors:** Rina Li, Xingcheng Lin

**Affiliations:** Department of Physics, North Carolina State University, Raleigh, NC; Bioinformatics Research Center, North Carolina State University, Raleigh, NC

## Abstract

Histone acetylation is a key regulatory post-translational modification closely associated with gene transcription. In particular, H4K16 acetylation (H4K16ac) is a crucial gene activation marker that induces an open chromatin configuration. While previous studies have explored the effects of H4K16ac on nucleosome interactions, how this local modification affects higher-order chromatin organization remains unclear. To bridge the chemical modifications of these histone tail lysine residues to global chromatin structure, we utilized a residue-resolution coarse-grained chromatin model and enhanced sampling techniques to simulate charge-neutralization effects of histone acetylation on nucleosome stability, inter-nucleosome interactions, and higher-order chromatin structure. Our simulations reveal that H4K16ac stabilizes a single nucleosome due to the reduced entropic contribution of histone tails during DNA unwrapping. In addition, acetylation modestly weakens inter-nucleosome interactions by diminishing contacts between histone tails, DNA, and nucleosome acidic patches. These weakened interactions are amplified when nucleosomes are connected by linker DNA, whose fluctuation causes significant chromatin destacking and decompaction, exposing nucleosomes for transcriptional activity. Our findings suggest that the geometric constraint imposed by chromatin DNA plays a critical role in driving chromatin structural reorganization upon post-translational modifications.

## Introduction

Epigenetic modifications are essential for regulating gene expressions within an organism. These modifications activate or repress gene expression, thereby controlling the transition between active and repressive chromatin.^1–5^ At the molecular level, histone modifications of nucleosomes constitute a key component of epigenetic modifications. Various post-translational modifications (PTMs) of histone tails control their interactions with nucleosomal DNA, further altering chromatin accessibility across different genomic regions and guiding various DNA-templated processes.^2,6–9^ While decades of research have revealed global structural rearrangement of chromatin driven by overall modification states of histone tails,^10–13^ only recently did techniques advance to obtain the resolution to study the effects of these histone modifications at an atomistic resolution. ^14^

Electrostatic interactions guide histone-regulated chromatin organization. Histone tails contain positively charged residues at multiple locations, which closely interact with the negatively charged nucleosomal DNA, thereby controlling inter-nucleosomal interactions and higher-order chromatin organization.^11^ Consequently, charge-modulating histone modifications guide structural changes in chromatin by modulating histone-tail-DNA interactions. Among these modifications, acetylation, particularly of histone 4 (H4) tails, is one of the most extensively studied charge-neutralizing histone modifications.^11–13,15,16^ H4 tail acetylation has been shown to activate gene expression^17^ by destablizing chromatin folding^13^ and promoting chromatin deassociation.^6,18,19^ Nonetheless, detailed information into atomistic interactions underlying histone-tail-mediated chromatin organization remains elusive.

Computational models can examine the atomistic details of histone-tail-DNA interactions, elucidating the structural impact of individual residues and their role in driving transitions among chromatin states. Numerous studies have demonstrated the effects of H4 acetylation on nucleosomal stability,^15,16^ histone-DNA interactions, ^20^ inter-nucleosomal interactions,^21^ and higher-order chromatin structures.^22,23^ These works have revealed how charge neutralization of positive lysine residues, coupled with secondary structural change of histone 4 tails, synergistically modulates chromatin structures. Nonetheless, due to the large size of a chromatin system (around 10,000 residues), it is challenging to directly connect the effects of individual acetylation to changes in overall chromatin organization by simulation. A chemically correct residue-resolution model can simulate large chromatin complexes while maintaining precise control of individual amino acids and nucleotides, ^23–30^ allowing us to focus on the modulatory effect of specific acetylated residues on higher-order chromatin structures.

In this study, we used a residue resolution chromatin model simulated in an explicit ion environment^30^ (Figure 1) to investigate the effects of histone tail acetylation, with a focus on the charge neutralizing effect of H4K16 acetylation (H4K16ac), a key modification known to activate gene expression.^13,17^ Specifically, we examined the impact of acetylation on internucleosome interaction and its connection to global chromatin structural rearrangement. Somewhat counterintuitively, our findings indicate that H4K16ac increases nucleosome stability by reducing the entropic penalty associated with the release of acetylated H4 tails. Our simulations also reveal a slight reduction in the inter-nucleosomal interaction incurred by H4K16ac. However, these small changes in inter-nucleosome interactions are amplified when nucleosomes are connected by DNA to form chromatin, leading to significant destacking of nucleosomes within the higher-order chromatin structure, thereby increasing nucleosome accessibility. Detailed analysis shows that H4K16ac decreases cross-nucleosome interactions between H4 tails, DNA, and acidic patches, resulting in large variations in the angles between flanking linker DNAs. Therefore, H4 acetylation primarily controls gene expression by disrupting nucleosome rearrangement through the amplification of weakened inter-nucleosome interactions at the chromatin level. Finally, we discuss the implications of our findings for other histone modifications.

**Figure 1:**
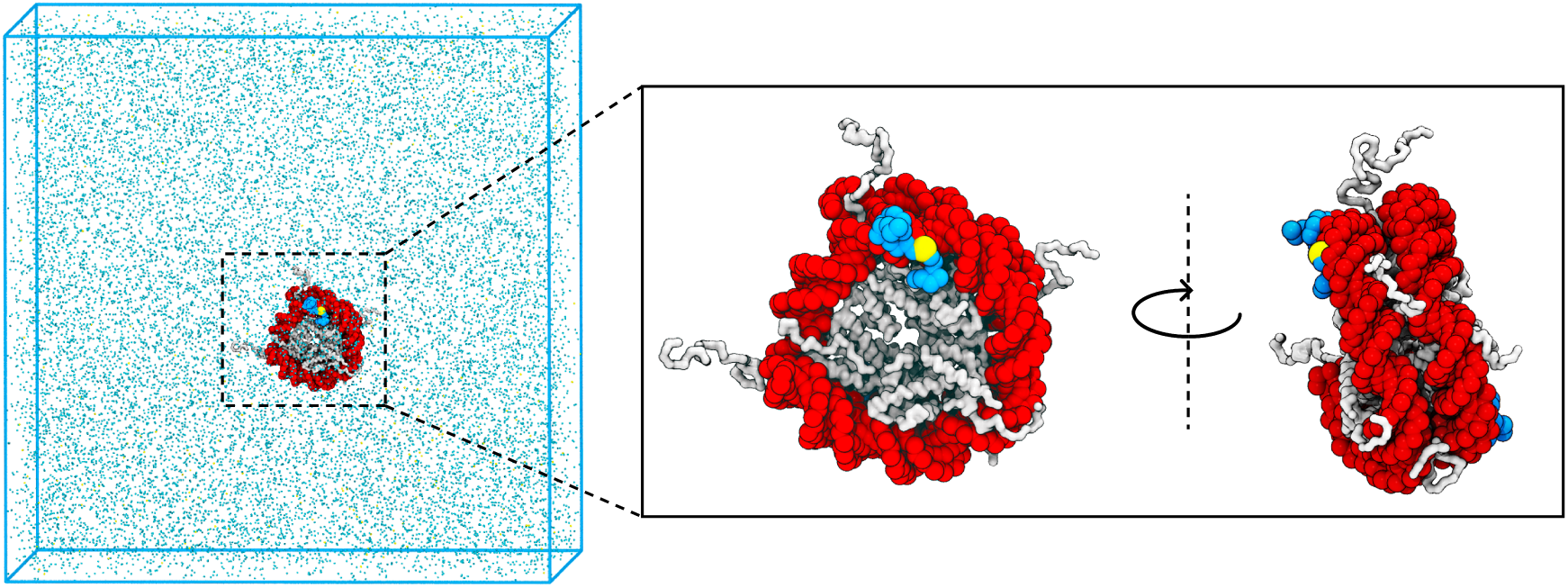
Illustration of the simulation model. The left panel displays the crystal structure of a single nucleosome, containing 147 base pairs of DNA within a simulation box, surrounded by explicitly modeled ions in physiological concentration (150 mM NaCl and 2 mM MgCl_2_). The nucleosomal DNA and histone proteins are colored red and white, respectively. The H4 tails are represented as blue spheres, with the Lysine 16 residue highlighted in yellow. The right panel provides a zoomed-in view, highlighting the relative position of the H4 tails and other nucleosomal components from two different perspectives.

## Results

### H4K16ac increases free energy penalty for nucleosomal DNA unwrapping

Numerous experiments have explored the energetic cost of chromatin unwrapping and its modulation by epigenetic modifications.^31–35^ Some studies directly probed the impact of histone tail acetylation on the unwrapping single nucleosomes by measuring the change of peak force needed to stretch nucleosomal arrays.^12^ Understanding the free energy penalty associated with unwrapping a single nucleosome can help disentangle contributions from intra- and inter-nucleosome interactions to chromatin organization.

We utilized a coarse-grained residue-resolution chromatin model (Figure 1) to simulate the unwrapping of DNA from a single nucleosome (see Section *Single nucleosome simulation* for details). The nucleosome was modeled with 147 base pairs of 601-sequence DNA and surrounded by explicit ions, similar to our previous work.^30^ The simulation was conducted in a physiological ion environment, including 150 mM NaCl and 2 mM MgCl_2_. Umbrella sampling^36^ was used to fully sample the unwrapping process of the nucleosomal DNA. To investigate the effect of acetylation, we simulated three types of nucleosomes: a wild-type 601-sequence nucleosome, one with the Lysine 16 neutralized on both the H4 tails to represent the chemical effect of H4K16ac marks, and another with all the Lysine residues on the H3 and H4 tails neutralized to represent the acetylation of both the histone tails.

Interestingly, the acetylated nucleosome shows a higher free energy cost for the unwrapping of the outer layer DNA, with an increase of 3.2 k*_B_*T in the barrier (Figure 2B). A detailed analysis of the energetics involved in the unwrapping process reveals the source of this increased free-energy penalty. Although neutralizing positive charges on the H4 tails lowers the energetic cost of unwrapping nucleosomal DNA, as indicated by the smaller reduction in contacts between the H4 tails and DNA (Figure 2C), the entropic cost associated with the H4 tails also decreases due to their increased conformational entropy after acetylation.^21^ The entropic contribution of histone tails to nucleosomal DNA unwrapping and histone-binding proteins interactions has been documented in earlier studies, ^28,37–39^ and our findings demonstrate that this effect can lead to a more stable nucleosome upon the introduction of H4K16ac. Acetylation of both H3 and H4 tails, however, would drastically remove the energetic contribution by histone-DNA interactions, significantly lowering the free energy cost of unwrapping the outer layer of nucleosome DNA, but not the inner layer. This is consistent with previous measurements from single-molecule optical trapping experiments.^12^

**Figure 2:**
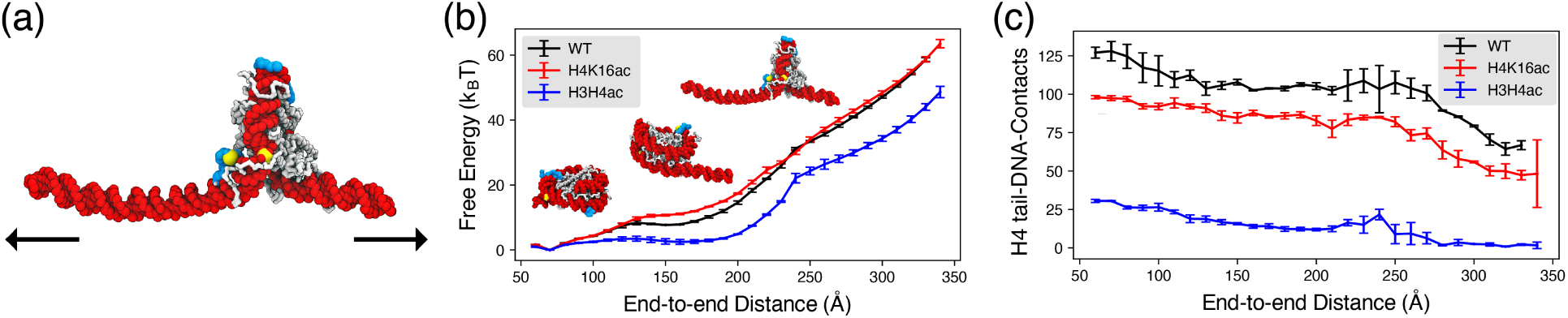
Acetylation changes the free energy penalty for unwrapping nucleosomal DNA. (a) Illustration of the umbrella sampling techniques used to unwrap a single nucleosome, with the end-to-end distance between the two DNA termini used as the umbrella collective variable. (b) Comparison of free energy profiles as a function of nucleosomal DNA end-to-end distance for the wild-type (WT, black), acetylated H4K16 (H4K16ac, red), and acetylated H3/H4 (H3H4ac, blue) nucleosomes. Error bars were calculated as the standard deviation of three independent estimates. (c) Comparison of H4 tail-DNA contact numbers as a function of nucleosomal DNA end-to-end distance for wild-type (WT, black), acetylated H4K16 (H4K16ac, red), and acetylated H3/H4 (H3H4ac, blue) nucleosomes. Error bars were calculated as the standard deviation of three independent estimates.

### Acetylated histone tails reduce inter-nucleosome interactions via weakened interaction with DNA and acidic patches

A more stable nucleosome cannot explain the experimental findings that showed chromatin unfolding upon the introduction of H4K16ac.^13^ To examine the source of this acetylation-induced chromatin unfolding, we next examine the effects of histone tail acetylations on inter-nucleosome interactions. A series of umbrella sampling simulations were performed to assess the inter-nucleosome interaction between two 601-sequence nucleosomes under the physiological salt concentration (150 mM NaCl and 2 mM MgCl_2_, Figure 3A, see Section *Two-nucleosome simulation* for details).

**Figure 3:**
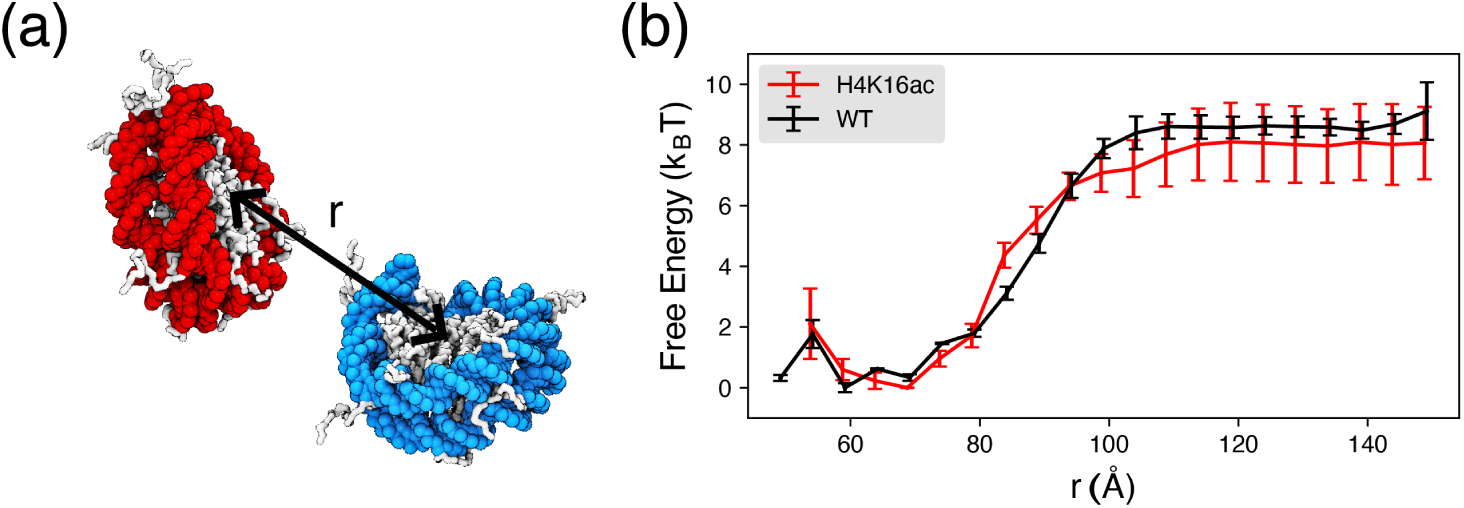
The effect of H4K16ac on inter-nucleosome binding. (a) Illustration of the inter-nucleosome distance r, measured between the geometric centers of two nucleosomes. (b) Free energy profile as a function of r, comparing two wild-type (WT) nucleosomes with two nucleosomes marked by H4K16ac modifications. Error bars were calculated as the standard deviation of three independent estimates.

Our simulation reveals a modest reduction in the inter-nucleosome interactions caused by H4K16ac modification (Figure 3B). The acetylation of H4 tails decreases the relative stability of the face-to-face binding mode between two nucleosomes compared to other binding configurations (Figure S1C), resulting in increased flexibility for nucleosomes to explore different conformations.

To examine the source of these weakened interactions, we calculated the average cross-nucleosome contacts between the H4 tails and nucleosomal DNA, as well as their interactions with the acidic patches of nucleosomes, a negatively charged area within the H2A-H2B histones (See Section *Details of simulation analysis* for details). Notably, the H4 tail forms long-range cross-nucleosome interactions with DNA, which are significantly weakened by H4K16ac modification (Figure 4A). Additionally, H4K16ac reduces the short-range interactions between H4 tails and the acidic patches. Therefore, both histone-tail-DNA and histone-tail-acidic patch interactions are weakened by the H4K16ac marks, thereby diminishing the inter-nucleosome interactions in the acetylated state.

**Figure 4:**
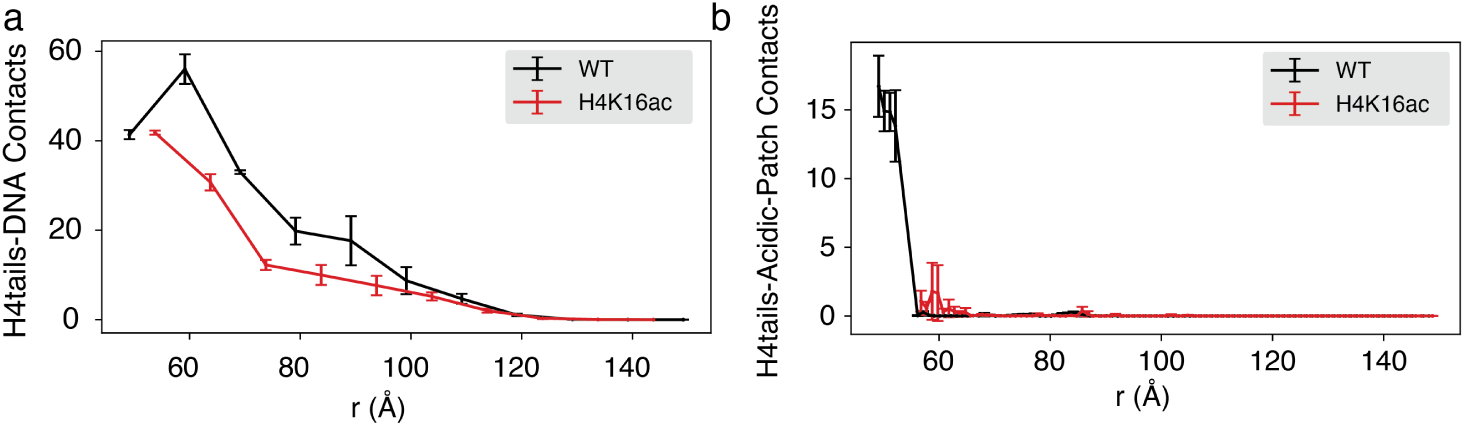
Acetylated H4 tails form fewer cross-nucleosome contacts with nucleosomal DNA and acidic patches. (a) Average cross-nucleosome contacts between H4 tails and nucleosomal DNA, compared between wild-type (WT) nucleosomes and those with H4K16ac modifications. Error bars were calculated as the standard deviation of the three independent estimates. (b) Average cross-nucleosome contacts between H4 tails and the acidic patches of nucleosomes, compared between wild-type (WT) nucleosomes and those with H4K16ac modifications. Error bars were calculated as the standard deviation of the three independent estimates.

### H4K16 acetylation induces significant chromatin destacking

Despite the weakened interaction between two nucleosomes, the acetylation-induced decrease in binding energy remains relatively minor compared to the extensive chromatin unfolding observed experimentally when introducing H4K16ac modifications.^13^ To further investigate the effect of H4K16 acetylation on the higher-order chromatin structure, we incorporated the H4K16 acetylated nucleosomes in a 12-nucleosomal (12-mer) chromatin array and carried out umbrella simulations^36^ to fully sample its global conformation (See Section *12-mer nucleosomal array simulation* for details). We used a collective variable, *Q*, which quantifies the deviation of a given 12-mer structure from a reference 12-mer two-start helical structure, with *Q* = 1.0 representing a perfect two-start compact structure and *Q* = 0.0 representing a completely disordered or extended chromatin structure.

As shown in Figure 5A, H4K16ac significantly destablizes the higher-order structure of the 12-mer chromatin, with the most populated structure featuring *Q* = 0.3. In contrast, the wild-type chromatin structure favors a more compact chromatin structure with *Q* between 0.5 and 0.7. To further understand the impact of H4K16ac on chromatin structure, we clustered and extracted the most populated structures from our simulation. Acetylation of Lysine 16 disrupts nucleosome-nucleosome stacking, exposing the nucleosomal DNA and increasing chromatin accessibility (Figure 5B).

**Figure 5:**
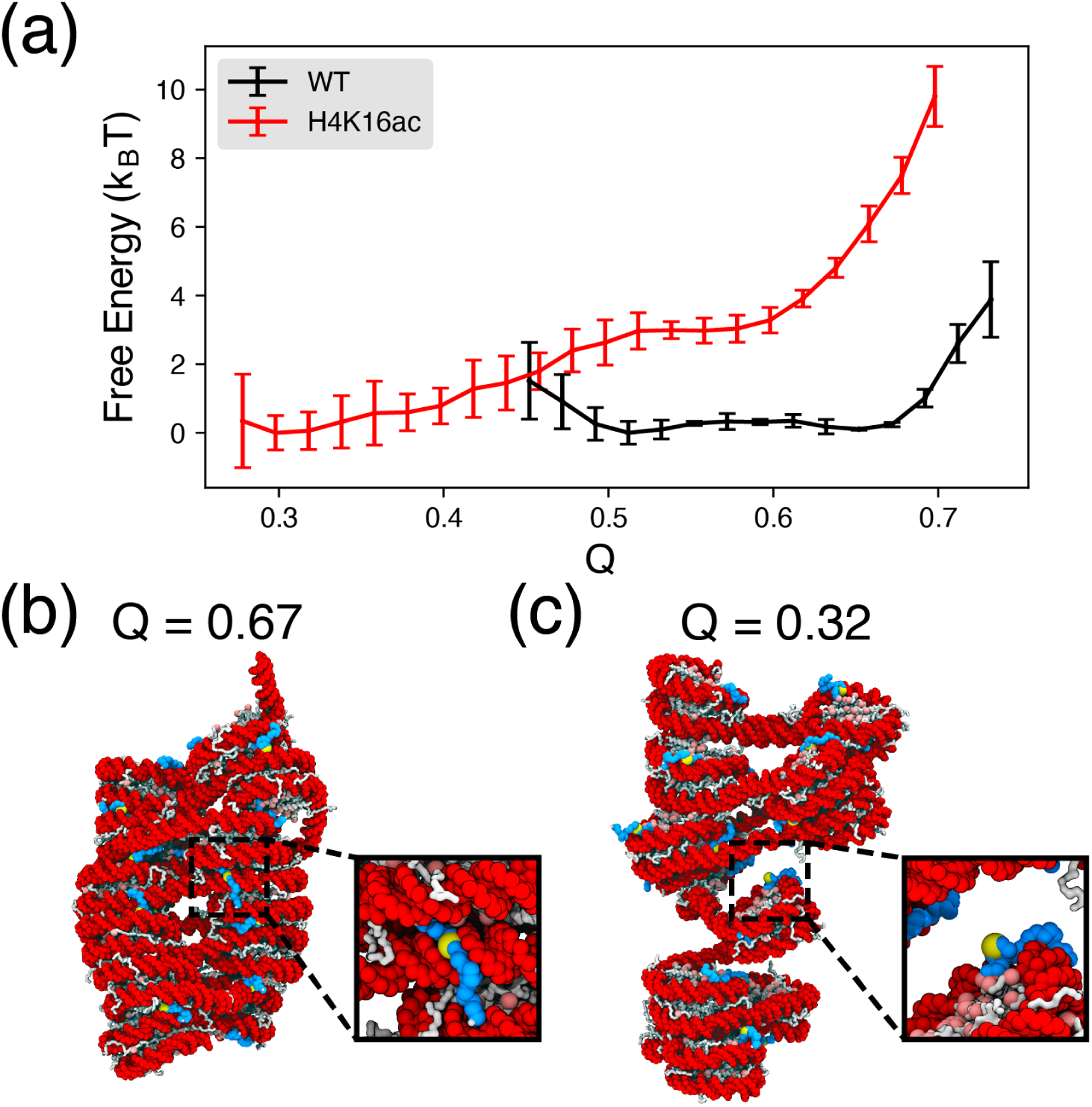
H4K16ac modification destabilizes the 12-mer chromatin structure. (a) Free energy profile as a function of *Q*, comparing wild-type and H4K16ac modified 12-mer nucleosomal arrays. The H4K16ac modification significantly destabilizes the chromatin structure, leading to a destacked conformation with a free energy minimum at *Q* = 0.3. (b) Representative chromatin structure of the wild-type 12-mer at its free energy minimum, with the corresponding *Q* value annotated. (C) Representative chromatin structure of the H4K16ac modified 12-mer at the free energy minimum, with the corresponding *Q* value annotated.

### Nucleosome linkage amplifies H4K16ac-weakened chromatin interactions

The observed destacking of chromatin is consistent with the experimental findings that H4K16ac modification induces chromatin unfolding and exposes DNA regions for gene activation.^13^ To further investigate the underlying mechanism of this structural decompaction, we performed a free-energy analysis to identify the key factors contributing to the acetylation-modulated chromatin structure.

Since chromatin compactness is primarily controlled by histone tail-nucleosomal DNA interactions, we computed the number of contacts between H4 tails and nucleosomal DNA in our 12-mer simulations. Consistent with our two nucleosome simulations (Figure 4), the contact number between the H4 tail and DNA decreases in the H4K16ac 12-mer array. This reduction in H4 tail-DNA interactions correlates with smaller *Q* values for the acetylated chromatin (Figure 6 a). In addition, since the entry-exit angle Ψ between linker DNA is directly linked to the higher-order chromatin structure,^40–43^ we computed the distribution of the average angle ⟨Ψ⟩ in our simulation (see Section *Linker DNA entry-exit angle* for details). The results show a greater fluctuation of ⟨Ψ⟩ of the H4K16ac modified chromatin (Figure 6 b). Therefore, the weakened inter-nucleosome interactions caused by H4K16ac modification are amplified when nucleosomes are connected by linker DNA, leading to an increased fluctuation of nucleosomal DNA at the entry-exit sites, contributing to the destacked chromatin structure.

**Figure 6:**
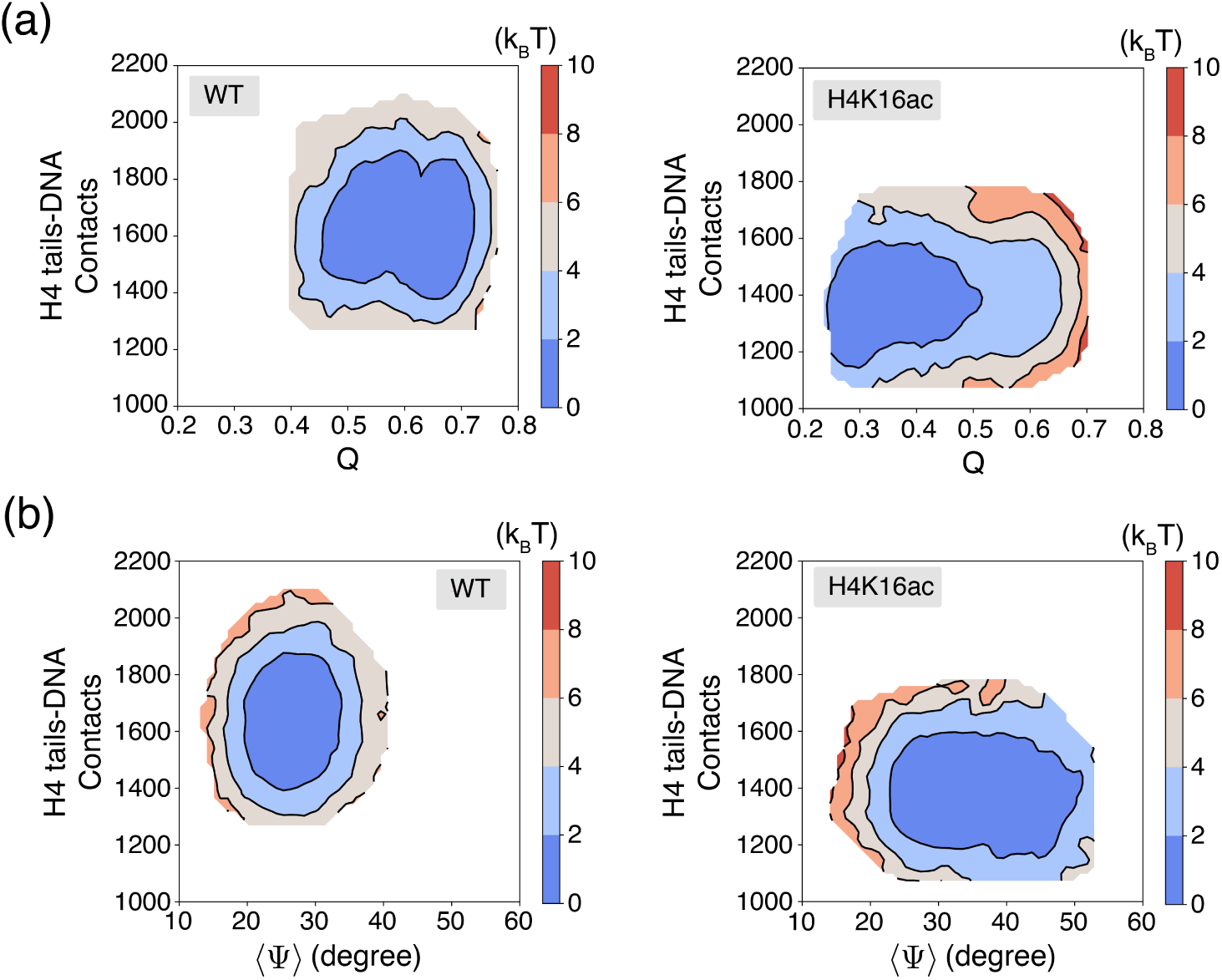
Reduced H4 tail-DNA contacts lead to a large fluctuation of linker-DNA entry-exit angles in acetylated chromatin, contributing to a destacked chromatin structure. (a) 2D free energy profile as a function of *Q* and the number of H4 tail-nucleosomal DNA contacts for the wild-type (WT) (left) and H4K16ac-modified (right) 12-mers. The H4K16ac-modified chromatin features fewer H4 tail-DNA contacts and a lower *Q* value. (b) 2D free energy profile as a function of the number of H4 tail-nucleosomal DNA contacts and Ψ, the average linker DNA entry-exist angle, for the wild-type (WT) (left) and H4K16ac-modified (right) 12-mers. The H4K16ac modified chromatin displays a larger fluctuation in ⟨Ψ⟩.

## Discussion

### Inter-nucleosome interactions as the primary source of weakened H4K16ac chromatin interactions

We utilized an explicit-ion, residue-resolution chromatin model to quantitatively characterize H4-tail acetylation and its modulation of chromatin interactions and organization. By investigating the effects of H4K16ac modification at different levels – within a single nucleosome, between two nucleosomes, and towards higher-order chromatin structures – we were able to disentangle the influence of H4K16ac on intra-nucleosome interactions from its impact on inter-nucleosome interactions. Interestingly, we found that the charge neutralization of H4K16 stabilizes nucleosomes by reducing the entropic force to release H4 tails from nucleosomal DNA. Acetylation also decreases cross-nucleosome interactions between H4 tails, DNA, and acidic patches, leading to a modest weakening of inter-nucleosome interactions. These weakened interactions are amplified at the chromatin level, leading to larger fluctuations in linker DNA angles, ultimately leading to significant chromatin destacking that exposes DNA for gene activation.

The reduced inter-nucleosome interactions also affect the phase-separation behaviors of chromatin. Inter-chain interactions between chromatin regions of the genome can stabilize the unfolded chromatin via interdigitation,^44–46^ resulting in a gel-like state with arrested kinetics in differentiated cells.^29,47,48^ H4K16ac weakens these inter-chain chromatin interactions, lowering the likelihood of chromatin association, a phenomenon proved by a previous precipitation experiment.^19^ Therefore, our simulation suggests that H4K16ac acetylation reduces the propensity for chromatin compartmentalization by phase separating into condensates, consistent with previous experimental findings.^49–51^

### Connected chromatin amplifies inter-nucleosome interactions through linker DNA

The compactness of chromatin determines the mechanical properties and transcriptional activities of genome.^50,52^ It is affected by various factors, including intrinsic physicochemical interactions within nucleosomes, such as histone-DNA and histone-acidic patch interactions, as well as extrinsic interactions with numerous chromatin regulatory complexes. ^35,53–56^ Notably, the entry-exit angles of nucleosomal DNA and the relative rotational angles between neighboring nucleosomes have been suggested to determine different chromatin conformations, which is supported by experimental observations.^40,57,58^ This hypothesis has been recently confirmed by cryogenic electron tomography (cryo-ET) *in vivo* experiment followed by statistical analysis.^43^ Our computational studies reinforce this statement by showing that the geometric connection of nucleosomes via linker DNA amplifies the weakened internucleosome interactions caused by H4K16ac, leading to greater variations of linker DNA angles (Figure 6) and ultimately contributing to chromatin destacking (Figure 5).

### Difference between H4K16ac and other histone post-translational modifications

Epigenetic modifications regulate gene activation and repression by introducing chemical modifications to histone proteins and DNA. ^3,59^ Beyond acetylation, hundreds of histone post-translational modifications have been identified in nucleosome, significantly influencing gene activities by altering molecular interactions that govern chromatin organization. ^9,60^ These modifications collectively form a “histone code” that dictates various gene expressions across different cellular stages.^61^ In addition to H4K16ac, acetylation of other histone tails, such as H2A/B and H3 tails, also modulates intra and inter-nucleosome interactions by modifying the histone tail rigidity and histone-DNA interactions. ^12,62^ The residue-resolution model is well-suited to study these histone modifications by adjusting the charge distribution and parameters of protein-DNA interactions. ^63–65^

## Methods and Materials

### Residue-resolution explicit-ion chromatin model

We employed a residue-resolution coarse-grained chromatin model with explicit treatment of ions based on their concentrations. Detailed descriptions of this model can be found in a previous publication.^30^ Specifically, the DNA was modeled with a coarse-grained 3SPN.2C model,^66^ and the proteins were modeled with a Cα-based structure-based model.^67,68^

Protein-protein interactions within histone proteins were modeled by the structure-based potential based on the initial configuration, generated with the Shadow algorithm^69^ using the SMOG software.^68^ Additionally, a residue-specific Miyazwa-Jernigan (MJ) potential^70^ was added to model the remaining protein-protein interactions, especially those between histones from different nucleosomes. We scaled the structure-based potential by 2.5 to avoid protein unfolding at 300K, and the MJ potential was scaled by 0.4 to provide a balanced treatment of both order and disordered proteins.^23^

The protein-DNA interactions were modeled with two potentials. A generic electrostatic Coulombic potential was used to model the electrostatic interactions, and a WCA potential was used to model the excluded volume effect. The Coulombic potential is defined as

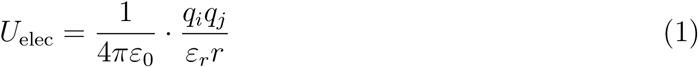

where ε_0_ is vacuum permittivity and ε*_r_* = 78.0 is the bulk solvent dielectric constant. *q_i_*, *q_j_* are the charges of two particles. The WCA potential is defined as a truncated and shifted Lennard-Jones potential at its minimum, with the expression as

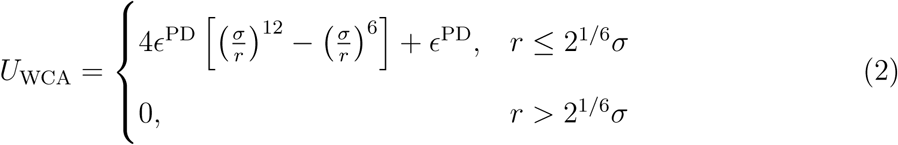

where σ = 5.7 Å and the interactive strength ɛ^PD^ is 0.02987572 kcal/mol.

The DNA-DNA interactions were modeled following a previous work, ^71^ with the parameters updated to the latest version of the DNA model, 3SPN.2C.^66^

### Coarse-grained explicit ion model

In our simulations, ions were explicitly modeled, with their interactions with protein and DNA described following previous works.^30,71^ Specifically, three components were used for the ion interaction potentials: an electrostatic potential described using the Coulombic form, a Gaussian potential to capture the solvent hydration effects, and a Lennard-Jones potential to represent short-range interactions.

The electrostatic potential between ion pairs is given by

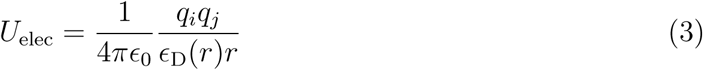

where *q_i_*, *q_j_* are the charges of two particles, and ɛ_0_ is the vacuum permmittivity, and ɛ_D_(*r*) is a distance-dependent dielectric constants. Specifically, the ɛ_D_(*r*) is calculated as follows:

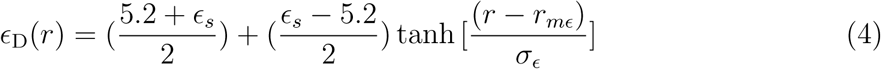

Here, ɛ*_s_*=78.0 is the dielectric constant of the bulk solvent, while *r_mɛ_* and σ*_ɛ_* represent the midpoint and the width of the transition between the saturated dielectric constant and the dielectric constant of the bulk solvent. The specific values of these constants are provided in Ref. 30.

The hydration potential follows a Gaussian form:

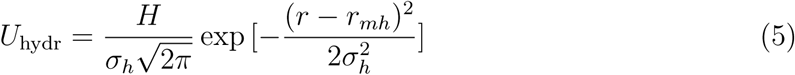

The hydration shell is characterized by three parameters: *r_mh_* for the midpoint, σ*_h_* for the width, and *H* for the height. The values of these ion-specific constants are available in Ref. 30.

The Lennard-Jones potential is described by:

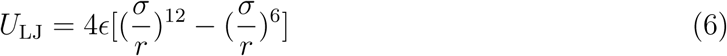

where the constants ɛ and σ are ion-specific constants, with their values listed in Ref. 30.

Total ion interaction is the sum of these three potentials

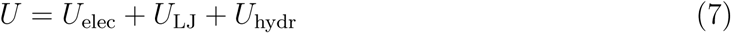

Further details about the parameters used in this model can be found in Ref. 71 and Ref. 30.

### Charge neutralization of acetylated histone tail sites

In all simulations, we modeled the acetylated lysine residues by neutralizing the positive charges of the corresponding residues in histone tails. Specifically, we modeled the H4K16ac acetyaltion by neutralizing Lysine 16 on all H4 tails. Additional ions were introduced to maintain charge neutrality and physiological ion concentrations across all systems.

### Details of simulation analysis

In this study, we conducted simulations on three systems: a single nucleosome, a two-nucleosome system, and a 12-mer nucleosomal array. The simulations were conducted using the software Lammps.^72^ To enhance the sampling of chromatin conformations and binding poses, we employed the umbrella sampling technique,^36^ which was implemented via the software PLUMED.^73^ We reweighted the simulation data to obtain the free energy by using the Weighted Histogram Analysis Method (WHAM),^74^ implemented through the SMOG software package.^68^

#### Single nucleosome simulation

We constructed a single nucleosome using the histone proteins from PDB ID 1KX5, replacing the 147-bp DNA sequence with the 601 sequence^75^ from the PDB structure 3LZ1.^76^ The ions were added and simulated in a cubic simulation box with an edge length of 600 Å, equilibrated with 150 mM NaCl and 2 mM MgCl_2_, along with additional ions to neutralize the system. Specifically, in the wild-type system, we used 19,129 Na^+^ ions, 260 Mg^2+^ ions, and 19,505 Cl*^−^* ions. For the H4K16ac system, 19,131 Na^+^ ions, 260 Mg^2+^ ions, and 19,505 Cl*^−^* ions were used, while the H3H4ac system included 19,155 Na^+^ ions, 260 Mg^2+^ ions, and 19,505 Cl*^−^* ions. All simulations were conducted at a temperature of 300 K.

To increase the computational efficiency, the histone core proteins and the two nucleotides on the dyad axis of the nucleosome were rigidified during the simulation. Umbrella sampling was performed using a collective variable (CV) defined as the end-to-end distance of the nucleosomal DNA, represented by the geometric center of the last five base pairs at each DNA terminus. During the simulation, we applied a spring constant of 0.001 kcal/(mol·Å^2^), with umbrella centers placed on a uniform grid from 30.0 Å to 510.0 Å at intervals of 30.0 Å, resulting in 17 umbrella windows. We simulated each umbrella trajectory for 13 million steps, with a time step of 10 fs, and the first 3 million steps were excluded to equilibrate the system when constructing the free energy profile.

#### Two-nucleosome simulation

To investigate nucleosome binding affinity and its modulation by H4K16ac, we simulated the binding free energy between two nucleosomes, each containing 147 base pairs of 601-sequence DNA, without the presence of linker histone or linker DNA. Each nucleosome was modeled as described in Section *Single nuclesome simulation*. To fully capture the binding free energy landscape between two nucleosomes, we performed umbrella sampling using two CVs (Figure S1 (a)). The first CV (depicted in Figure 3 (a) and S1 (a)), *r*, was defined as the distance between the geometric centers of two nucleosomes. Each nucleosome was represented by a list of residues: 63-120, 165-217, 263-324, 398-462, 550-607, 652-704, 750-811, 885-949, based on the residue IDs used in the nucleosome crystal structure (PDB ID: 1KX5).^26^ The second CV, θ, was defined as the angle between two unit vectors perpendicular to the nucleosome faces, w⃗_1_ and w⃗_2_.

Umbrella sampling was performed with a spring constant of 0.01 kcal/(mol·Å^2^) for r and 0.001 kcal/(mol·◦^2^) for θ. The histone core proteins and inner-layer DNA of each nucleosome were rigidified during the simulations to increase computational efficiency. Umbrella centers were placed on a uniform grid, with r ranging from 60.0 Å to 130.0 Å, and θ from 0*^◦^* to 180*^◦^*, with intervals of 10.0 Å and 30*^◦^*, resulting in 56 umbrella windows. Each umbrella trajectory was simulated for 12.75 million steps, with a timestep of 10 fs, and the first 3 million steps were excluded when constructing the unbiased binding free energy profile (Figure 3, S1).

The system was simulated in a cubic box of 600 Å, equilibrated with ions to ensure a physiological concentration of 150 mM NaCl and 2 mM MgCl_2_, with additional ions for neutralization. Specifically, the wild-type system contained 19,505 Na^+^, 260 Mg^2+^, and 19,737 Cl*^−^*, and the H4K16ac modified system had 19,509 Na^+^, 260 Mg^2+^, and 19,737 Cl*^−^*. All simulations were carried out at a constant temperature of 300 K.

#### 12-mer nucleosomal array simulation

Our simulation of the 12-mer nucleosomal array consists of 12 nucleosomes connected by 20 base pairs of linker DNA between neighboring nucleosomes. The construction of this array follows the protocol described in previous publications.^29,30^ Briefly, we used a single nucleosome and its associated linker DNA from the tetranucleosome crystal structure (PDB ID: 1ZBB)^77^ and connected 12 such units. The extra 20 base pairs of linker DNA from the last unit were removed to complete the system setup.

To fully sample the conformational space of this 12-mer chromatin, we used a CV, *Q*, for umbrella sampling. *Q* measured the structural deviation of the 12-mer array from a reference two-start compact chromatin structure, using the following equation:

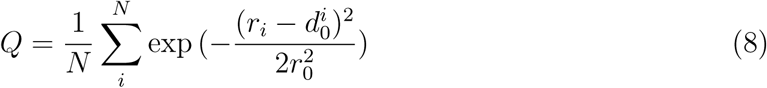

In this equation, *N* = 66 represents the total number of nucleosome pairs in the 12-mer array, *r_i_* is the distance between nucleosomes in the ith pair, r_0_ = 20.0 Å, d*^i^* denotes the distance between the ith pair of nucleosomes in the two-start reference structure. The value of *Q* ranges from 0 to 1, where larger values indicate higher similarity to the reference two-start chromatin structure, and thus representing a more compact chromatin. All the CVs used in our simulations were calculated using PLUMED.^73^

We performed umbrella sampling using *Q* as the CV with a spring constant of 50.0 kcal/mol. The histone core proteins and inner-layer DNA were rigidified during the simulations to increase computational efficiency. Six umbrella windows were simulated, with the umbrella centers placed on a uniform grid from 0.40 to 0.90 with an interval of 0.1. Each umbrella trajectory was simulated for 15 million steps, using a time step of 10 fs. The first 3 million steps were excluded from the free energy profile construction.

The 12-mer array was simulated in a cubic box of 600 Å, equilibrated with 150 mM NaCl and 2 mM MgCl_2_, along with additional ions to neutralize the system. In the wildtype 12-mer nucleosomal array simulation, we included 21,695 Na^+^ ions, 260 Mg^2+^ ions, and 20,025 Cl*^−^* ions. For the H4K16ac 12-mer nucleosomal array simulation, we included 21,719 Na^+^ ions, 260 Mg^2+^ ions, and 20,025 Cl*^−^* ions, with K16 neutralized on the H4 tails of all 12 nucleosomes. All simulations were carried out at a constant temperature of 300 K.

### Details of simulation analysis

This section describes the collective variables (CVs) used in our simulation analysis. The primary tools for the analysis were the PLUMED software package ^73^ and the Weighted Histogram Analysis Method (WHAM) implemented in the SMOG software package.^69^ PLUMED was used to compute key CVs, while WHAM was used to remove the biases introduced by the umbrella sampling potential.

#### Histone 4 tail-DNA contacts

We calculated the number of contacts between the H4 tails and DNA using PLUMED.^73^ For the 12-mer nucleosomal array, we also computed the contact numbers between the H4 tails and the linker DNA for comparative analysis. A higher contact number indicates stronger and more frequent interactions between these components.

To calculate the contact number, we used the COORDINATION command in PLUMED. The total number of contacts (*CN*) between the H4 tails and DNA was computed using the equation below:

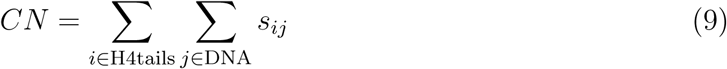

In this equation, *CN* represents the total number of contacts between H4 tails and DNA. The function s*_ij_* is a switching function that determines whether a contact is formed based on the distance between particles in the two groups:

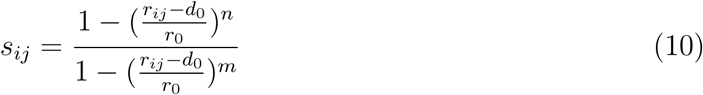

Here, *r_ij_* is the distance between particles in the two groups, *d*_0_ = 0.0, *r*_0_ = 8.0Å, *n* = 6, and *m* = 12.

#### Histone 4 tail-acidic patch contacts

Similar to the H4-DNA interaction analysis, we computed the number of contacts between the H4 tails and the acidic patches of nucleosomes using PLUMED, using the same functions and parameters. Following a previous study,^21^ we defined the acidic patch as consisting of the glutamic acid and aspartic acid residues, with residue ID: 56, 61, 64, 90, 91, 92, 102, and 110 on each H2A-H2B dimer.

#### Linker DNA entry-exit angle

For the 12-mer nucleosomal array system, we calculated Ψ, the angle between vectors representing the entry and exit linker DNAs for each nucleosome. Since B-form DNA has 10 base pairs per turn, the direction of each linker DNA was represented by a vector connecting the geometric center of the nucleosomal DNA entry/exit site to the geometric center of another base pair 10 base pairs away from the nucleosome (Figure S2). The angle between the two linker DNA was calculated using the following equation:

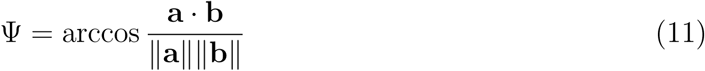

The average angle ⟨Ψ⟩ was then calculated as:

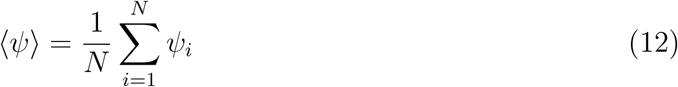

Since the first nucleosome lacks an entry linker DNA and the last nucleosome lacks an exit linker DNA, we averaged Ψ over the middle 10 nucleosomes in the 12-mer array (*N* = 10). All angle calculations were performed using PLUMED.

## Acknowledgement

This work was supported by the startup funding from North Carolina State University. This research was also funded, in part, by the NC State Genetics and Genomics Academy.

