## Supplementary Information for "Connected Chromatin Amplifies Acetylation-modulated Nucleosome Interactions"

### Supporting Information: Connected Chromatin Amplifies Acetylation-modulated Nucleosome Interactions

Rina Li[†] and Xingcheng Lin[*,†,‡]

#### Contents

**1 Figures** **S-2**

### 1 Figures

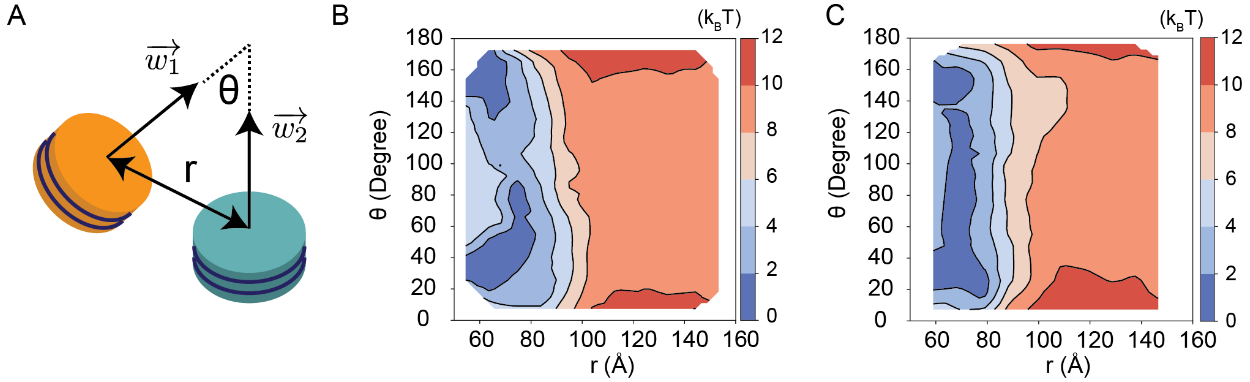


Figure S1: **Illustration of two collective variables used in umbrella sampling simulations of inter-nucleosome interactions, and the corresponding binding free energy profiles** (a) Schematic representations of nucleosomes showing $\vec{w_1}$ and $\vec{w_2}$, which are vectors perpendicular to the two nucleosomal faces. The first collective variable, $\theta$, represents the angle between these two vectors and thus the angle between the two nucleosomal faces. The second collective variable, $r$, represents the distance between the geometric centers of the two nucleosomes. (b) The 2D binding free energy profile between two wild-type 601-sequence nucleosomes, as a function of $\theta$ and $r$, at physiological salt concentration (150 mM NaCl and 2 mM $MgCl_2$). (c) The 2D binding free energy profile between two H4K16ac modified 601-sequence nucleosomes, as a function of $\theta$ and $r$, at physiological salt concentration (150 mM NaCl and 2 mM $MgCl_2$). Panel (A) and (B) are adapted from Ref. S1.

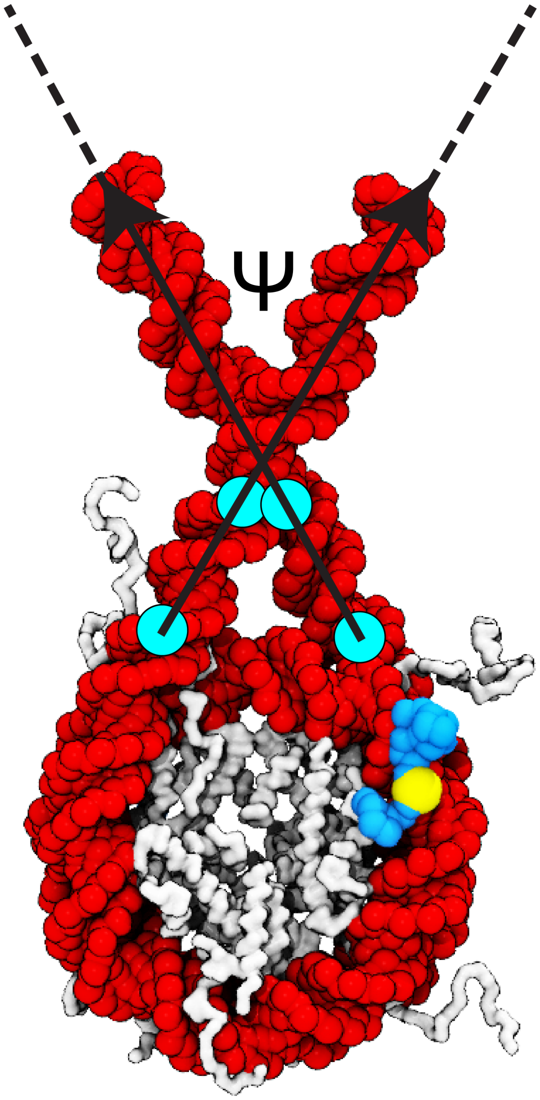


Figure S2: **Illustration of Ψ, the angle between entry and exit linker DNAs** The figure use one nucleosome to illustrate the calculation of Ψ, the angle between the entry and exit linker DNAs. The base pairs used to define Ψ are highlighted in cyan spheres. Ψ is defined as the angle between two arrows representing the directions of the entry and exit linker DNAs.

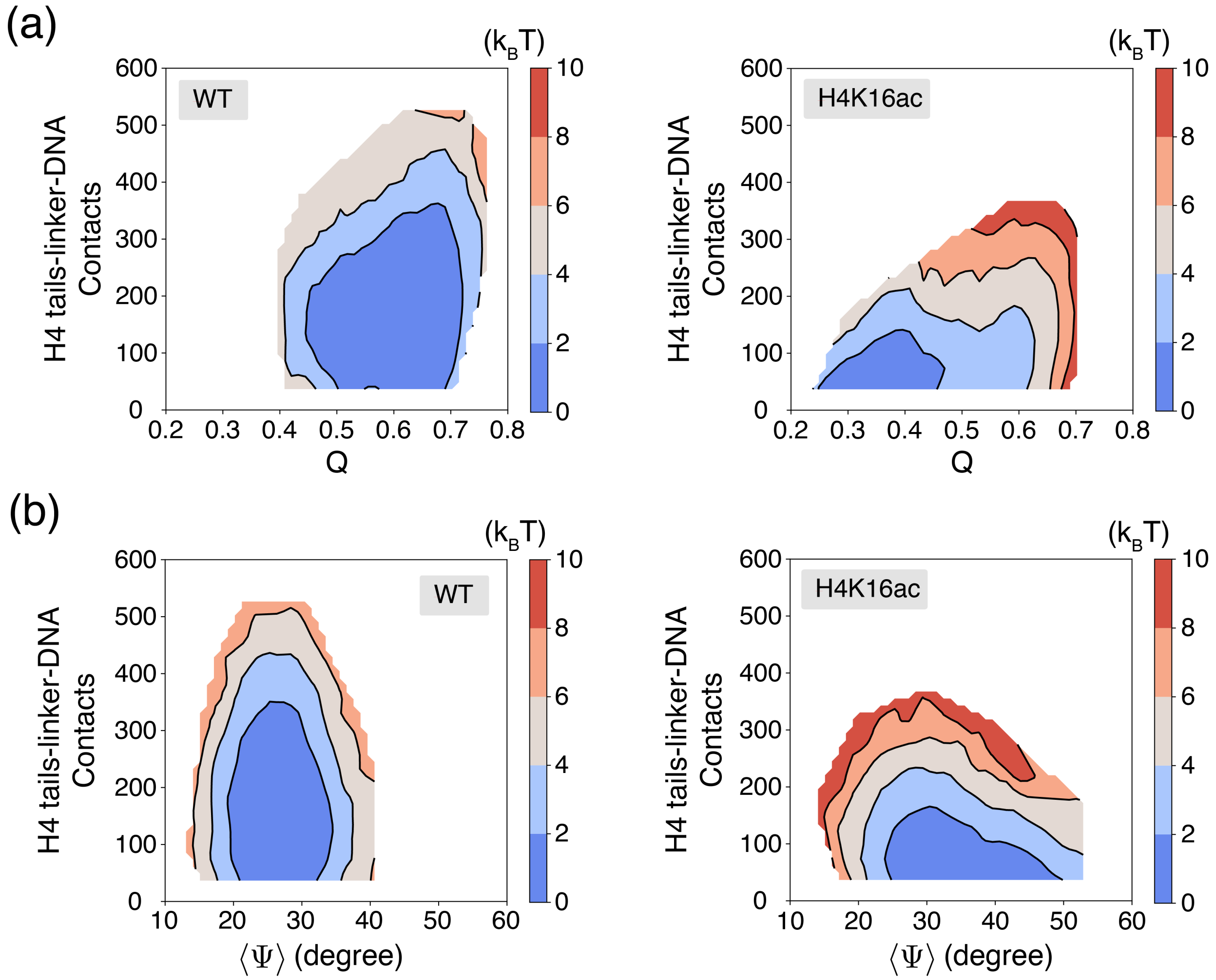


Figure S3: **Reduced H4 tail-DNA contacts lead to a large fluctuation of linker-DNA entry-exit angles in acetylated chromatin, contributing to a destacked chromatin structure, related to Figure 6** (a) 2D free energy profile as a function of $Q$ and the number of H4 tail-linker DNA contacts for the wild-type (WT) (left) and H4K16ac-modified (right) 12-mers. The H4K16ac-modified chromatin features fewer H4 tail-linker DNA contacts and a lower $Q$ value. (b) 2D free energy profile as a function of the number of H4 tail-linker DNA contacts and $\langle \Psi \rangle$, the average linker DNA entry-exist angle, for the wild-type (WT) (left) and H4K16ac-modified (right) 12-mers. The H4K16ac modified chromatin displays a larger fluctuation in $\langle \Psi \rangle$.

(S1) Lin, X.; Zhang, B. Explicit ion modeling predicts physicochemical interactions for chromatin organization. *eLife* **2024**, *12*, RP90073.
